# Discovery of a novel drug using lipid-based formulation targeting G12D-mutated KRAS4B through non-covalent bonds

**DOI:** 10.1101/2023.04.05.535682

**Authors:** Huixia Lu, Zheyao Hu, Jordi Faraudo, Jordi Martí

## Abstract

One of the most common drivers in human cancer is KRAS4B. In recent years, the promising KRAS targeted drug development has attracted significant new research interest and reignited the field of RAS therapeutics. To signal, oncogenic KRAS4B not only requires a sufficient nucleotide exchange, but also needs to recruit effectors by exposing its effector-binding sites while anchoring to plasma membrane where KRAS4B-mediated signaling events occur. Phosphodiesterase-*δ* plays an important role in sequestering KRAS4B from the cytoplasm and targeting it to cellular membranes. In this work, we have designed a drug LIG1 using lipid-based formulation to directly target both the switch-*II* pocket of KRAS4B-G12D and phosphodiesterase-*δ*. LIG1 was found to lock KRAS4B in its GDP-bound state while the effector-binding domain is blocked by the interface of the plasma membrane which hinders the nucleotide exchange while simultaneously it can affect the GTP-bound KRAS4B to shift from an active state to its inactive state. LIG1 is also observed to stably accommodate itself in the prenyl-binding pocket of phosphodiesterase-*δ* which impairs KRAS4B enrichment at the membrane and suppress the proliferation of KRAS4B-dependent cancer cells. In this work we report a drug based on lipid-based formulation that can foster drug discovery efforts for the targeting of oncogenes of the RAS family and beyond.

## Introduction

RAS proteins are small, membrane-anchoring guanosine-triphosphatase (GTPase) species that, in order to regulate cell survival and proliferation, oscillate between inactive GDP-bound and active GTP-bound states.^1^ They work as GDP-GTP binary switches and regulate cytoplasmic signalling networks able to control cellular processes involved in cell behaviour so that overacting RAS signalling can lead to cancer. RAS plays a crucial role in the regulation of different signaling pathways in cells. Proteins adopt unstable, high-energy states that exist for fractions of a second but can have key biological roles. ^2^ This is normally due to GTPase activating proteins such as neurofibromin or by activation of guanine exchange factors such as human regulator of chromosome condensation. The major RAS isoforms are encoded by three genes, resulting in a total of four RAS proteins (KRAS4A, KRAS4B, HRAS, and NRAS). Interest has largely been given to the oncogenic mutants of KRAS4B due to its relatively high frequency in cancers, in particular in 17% of lung, 61% of pancreas, 20% of small intestine, 33% of large intestine, and biliary tract cancer cells.^3–6^ Whereas the classic definition of an active RAS state is based on the GTP/GDP binding status, conformational reasoning argues that on its own, GTP-bound RAS may not be sufficient for the downstream effector and upstream regulator interactions while RAS proteins associating with the cell-specific plasma membrane (PM) bilayers, where the RAS signaling initiate.^7^

An analysis of mutation patterns across the RAS isoforms reveals that 66% of KRAS mutations occur at codon 12. Notably, at codon 12, KRAS is predominantly G12D mutated. ^4^ The high frequency of this mutation makes it an ideal drug target. However, attempts to target mutant RAS proteins have proven challenging.^8, 9^

The catalytic domain (CD, residues 1-166) of KRAS4B is composed of the effector-binding domain (ED, residues 1-86) and the allosteric domain (AD, residues 87-166). Four regions border the nucleotide-binding pocket: the phosphate-binding loop (P-loop, residues 10-17), switch *I* (S*I*, residues 30-38), switch *II* (S*II*, residues 60-76), and the base binding loops (residues 116-120 and residues 145-147).^10^ KRAS is distinguished by their C-terminal hypervariable region (HVR, res 167-185) whereas KRAS4B can go through post-translational modifications. HVR preferentially binds the membrane in the liquid phase and spontaneously inserts its terminal farnesyl moiety (FAR) into the loosely packed phospholipid/cholesterol bilayers of the main structure of the PM.^6^

From previous studies carried out in our lab,^11, 12^ the conformational behavior of KRAS4B-G12D at the anionic model PM has been systematically explored and the corresponding conformational landscape of two defined torsion angles has been revealed by using a joint method of molecular dynamics (MD) and well-tempered metadynamics simulations, which is of importance in modulating specificity. MD is a powerful tool which allows us to monitor the trajectory of each individual atom in the system in a wide variety of setups, ranging from simple liquids to biological membranes. MD can, in addition, obtain energetic, structural and especially time-dependent properties such as the diffusion coefficients or spectral densities,^13, 14^ enhancing its applicability. To signal, KRAS4B that is delocalized on endomembranes requires anchorage into PM of different local compositions. ^15^ The content of PM is reported to regulate the localization and activity of KRAS4B,^16^ favoring acidic disordered membranes where phosphatidylinositols, phosphatidylserine, and some other microdomain components are involved. The main interest of the oncogenic KRAS4B relies on its affinity to the PM and its enhanced protein recruitment as revealed by several works.^17–20^

To date, different small-molecule inhibitors of RAS proteins have been reported by different groups, although their potency and specificity are still limited.^10, 21–25^ A report from clinical evidence on the druggability of KRAS has shed some light on this long-term standing question.^26^ While many researchers have studied and provided insights into the interactions between small-molecules and different KRAS4B variants, further investigation is needed to explore different KRAS4B conformations aimed by new possible anti-cancer drugs for better understanding the mechanisms of KRAS4B-G12D driven cancer research. Recently, the regulation of oncogenic KRAS4B activity when binding to the cell membrane has been thoroughly investigated.^12, 27^ A crucial point is to locate druggable pockets at the surface of the protein. Some authors have reported indications of those allosteric pockets that may allow small-molecules to target these particular sites. ^28–31^

Currently, selective targeting of RAS in the clinic is limited to KRAS (G12C mutant) through covalent inhibitors (AMG510^32^ and MRTX849^33^). These two drugs have reached a preliminary, yet important success targeting KRAS G12C mutant. Thus, covalent inhibitors with a covalent moiety (warhead)^34^ show great potential as anticancer drugs. An electrophilic group, the warhead, of a covalent inhibitor can form covalent bonds with amino acids such as CYS, LYS, or TYR on targeted proteins.^35^ Examples of warheads adopted in the drug design are shown in Figure S1 (see Supporting Information). The affinity of the warhead to the target is of great importance for developing effective inhibitors.

Two major drug-binding paradigms (induced-fit and conformational selection) have superseded the static lock-and-key binding paradigm. ^36, 37^ Advanced NMR and single-molecule experiments indicate that the conformational change of a protein occurs prior to a binding event (’conformational selection’), rather than after a binding event (‘induced fit’). ^38^ MD methods are well suited for drug design, since they are based on equilibrated conformations of the target protein evolving along simulatime, where the flexibility of its structure can be explored during long MD trajectories, in comparison to other methods like docking in which only static structures are considered.

3,4-dihydro-1,2,4-benzothiadiazine-1,1-dioxide (C_7_H_8_N_2_O_2_S, DBD) is a bicyclic heterocyclic benzene derivative featuring amphiphilic properties and providing an opportunity as pharmacophores/warheads during the process of drug design. Its derivatives^39, 40^ have been long used in the human therapy as diuretic and antihypertensive agents, due to their widely biological pharmacological activities.^41, 42^ Lipid-based drug delivery systems, one of the most studied bioavailability enhancement technologies, are utilized in a number of U.S. Food and Drug Administration (FDA) approved drugs,^43^ some chemical structures of some FDA approved drugs shown in Figure S2. A considerable percentage of highly potent drugs are both poorly water-soluble and poorly permeable which may lead to translates into suboptimal patient outcomes due to poor oral bioavailability and variable pharmacokinetics.^44^ The overall drugs based on lipid-based formulation overall have very good permeability properties in human jejunum.^43^

Active KRAS signalling occurs at the PM membrane. And increasing numbers of ligand-bound membrane protein structures challenge traditional notions of drug activity derived from ligand–protein interactions. ^45^ It suggests that interactions of ligand-membrane, ligand-target protein at the presence of membranes with different compositions are likely underappreciated components of drug discovery. PM interacting with drug molecules is not a new concept. However, the implications of PM to a drug receptor such as KRAS mutants are hardly addressed in the modern medicinal chemistry literature. We believe PM plays a key role in the process of different drug pathways, thus requires deeper consideration in light of the development of anti-carcinogens or tumour-reducing drugs.

In this work, we address a question of both fundamental and practical importance about how a lipid-based drug (LIG1) may provide a novel strategy to reduce oncogenic KRAS4B-G12D mutant signaling. By applying MD simulations in the microsecond time scale, we gain insight into the role of LIG1 and DBD drug molecules might play for KRAS4B-G12D in its GTP-bound and GDP-bound states. Exploring whether LIG1 can non-covalently inhibit KRAS4B-G12D, other RAS isoforms and their mutants will give wider application to different RAS-driven cancers. And the lipid-based formulation may to be applicable to other important therapeutic targets and presents the exciting potential for an alternative pipeline for drug discovery. Our studies will show that LIG1 can inhibit KRAS4B-G12D’s signaling pathways by two means (Fig. 1), as it will be described in the following sections..

**Figure 1:**
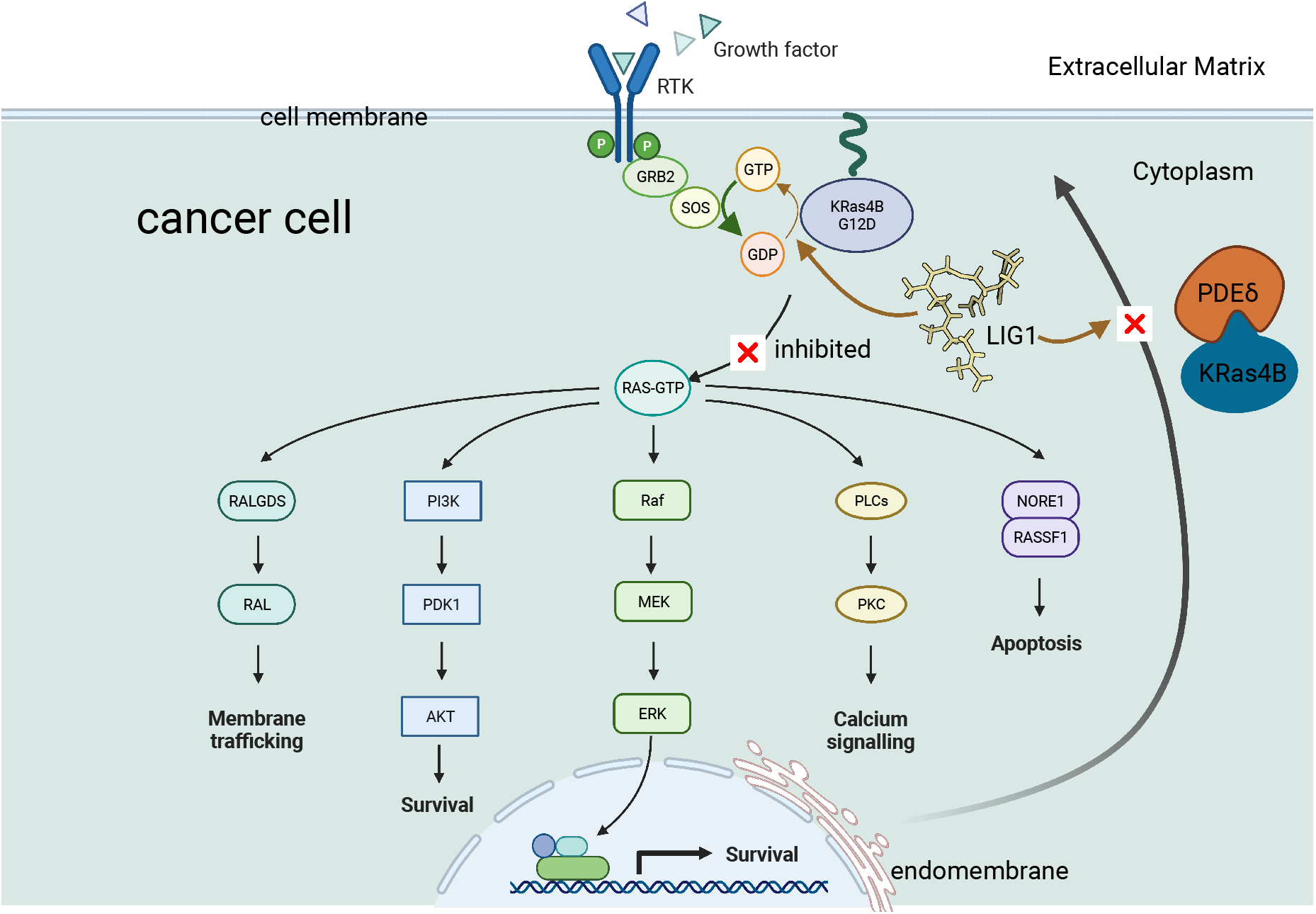
LIG1 blocks KRAS4B signaling pathways through two means, resulting KRAS4B in its GDP-bound inactive states and an impaired enrichment of KRAS4B at the inner leaflet of PM. Figure adapted from “Ras Pathway” by BioRender.com (2023).

## Results and Discussion

In this work we have performed MD simulations at the microsecond scale (in total 15.81 µs) in different situations to investigate the role of the plasma membrane, LIG1, and DBD play in the dynamics behavior of KRAS4B-G12D. Details of designed MD simulations are summarized in Table 1.

**Table 1:**
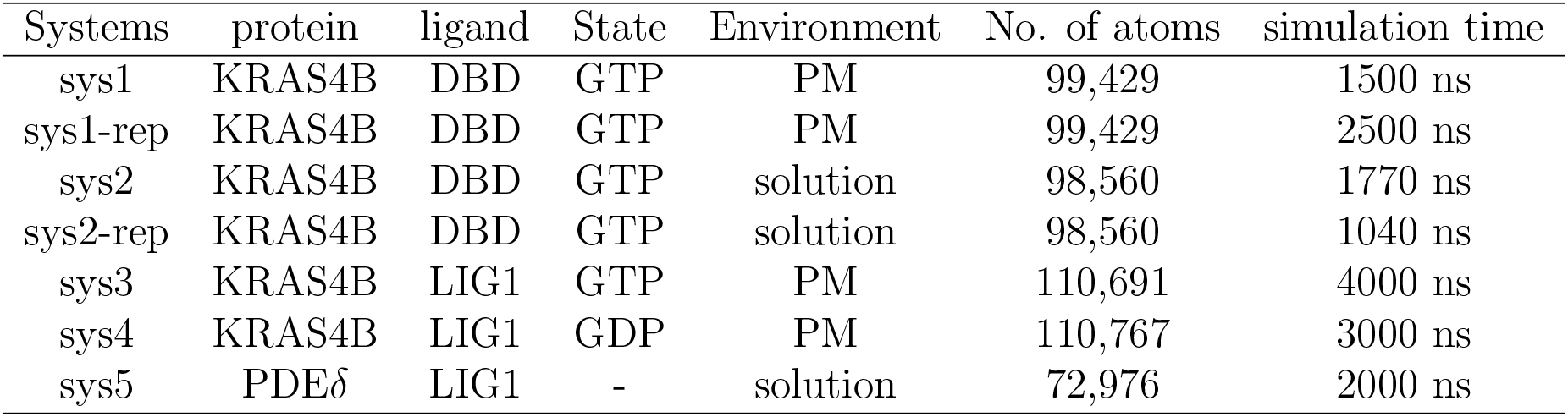
Detailed setups of all simulation systems considered in this work. Among them, sys1-rep and sys2-rep which stand for independent production runs for the same system, more details can be found in Supporting Information (SI).

| Systems | protein | ligand | State | Environment | No. of atoms | simulation time |
| --- | --- | --- | --- | --- | --- | --- |
| sys1 | KRAS4B | DBD | GTP | PM | 99,429 | 1500 ns |
| sys1-rep | KRAS4B | DBD | GTP | PM | 99,429 | 2500 ns |
| sys2 | KRAS4B | DBD | GTP | solution | 98,560 | 1770 ns |
| sys2-rep | KRAS4B | DBD | GTP | solution | 98,560 | 1040 ns |
| sys3 | KRAS4B | LIG1 | GTP | PM | 110,691 | 4000 ns |
| sys4 | KRAS4B | LIG1 | GDP | PM | 110,767 | 3000 ns |
| sys5 | PDEδ | LIG1 | - | solution | 72,976 | 2000 ns |

We have observed that FAR is able to bind stably into S*II* pocket of KRASB-G12D in aqueous environment for 1.5 *µ*s of simulation time, obtained by distance of center of mass of FAR and CD of Figure S3 B, showing its potential act as the anchor to target KRAS4B mutants. Fig. 2 shows how we designed LIG1 by Chimera^46^ that composes a lipid tail (FAR domain) and a hydrophilic head (DBD domain) according to the lipid-based formulation as our results here suggest, allowing LIG1 to preserve properties of good permeability across PM.

**Figure 2:**
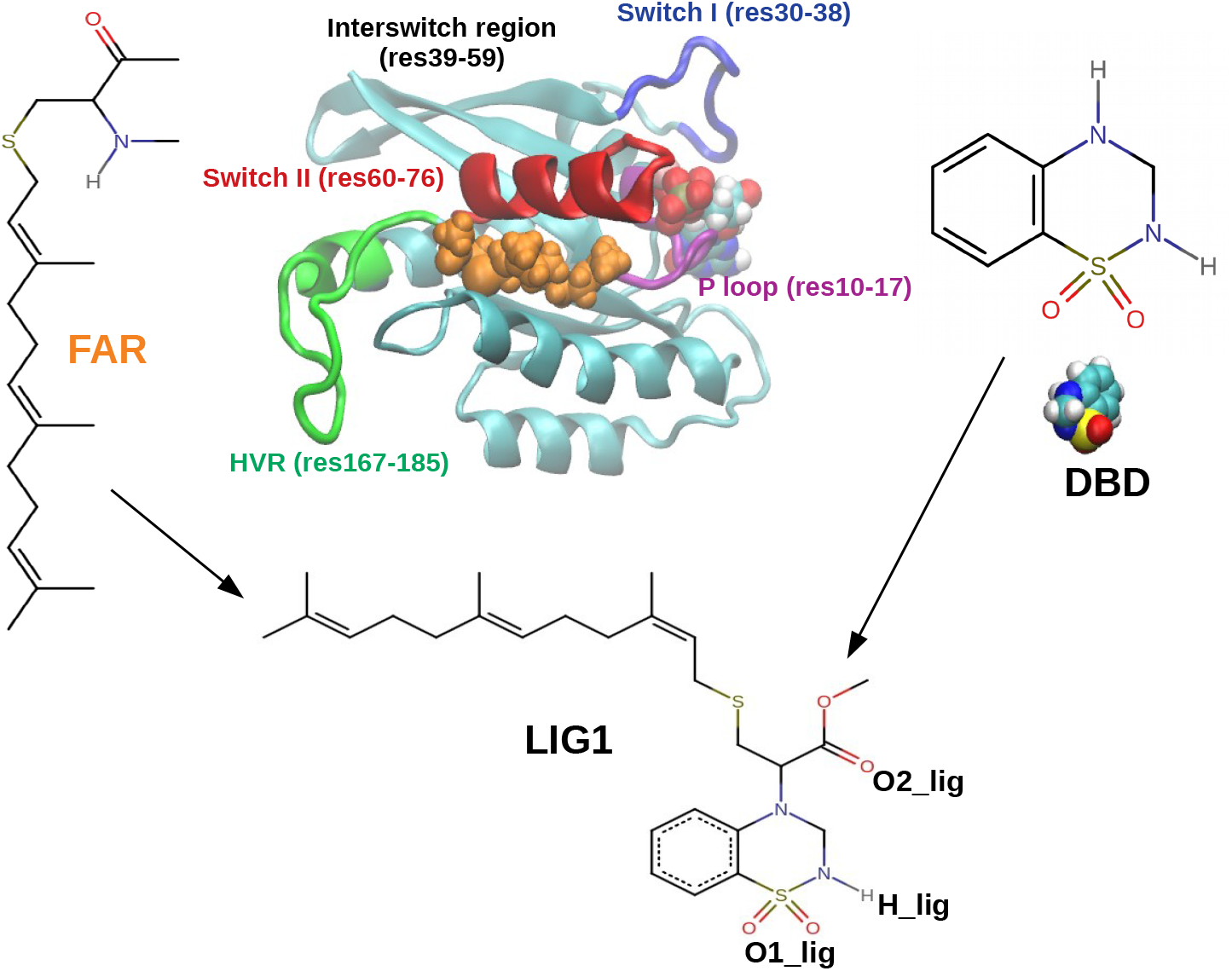
FAR binds to CD mainly by means of hydrophobic force with S*II*, together with the help of hydrogen bonds within residues of HVR. Chemical structures of FAR, DBD, LIG1 are shown here. In a closer look of KRAS4B of sys2, HVR, S*I*, S*II*, and P loop are shown in newCartoon representation in color green, blue, red, and purple, respectively. FAR that interacts with S*II* of KRAS4B solvated in solution is emphasized in orange Van Der Waals representation.

### Structural characteristics of the oncogenic KRAS4B

KRAS4B was considered as an undruggable protein, but now is rather considered as a challenging target. Despite the recent substantial progress in KRAS4B drug discovery, we still do not fully understand the underlying mechanisms of anticancer drug targeting KRAS4B at the presence of PM. According to structural data^47^ it may easily lead one to the false assumption: the switch regions stay in the stabilized conformations. It was believed that the two switch regions differ in conformation between the GDP and GTP states.^48^ THR35 and GLY60 were reported to make hydrogen bonds with the *γ*-phosphate, thus hold S*I* and S*II* regions in their active conformations in the GTP state. When GTP is hydrolyzed, S*I* and S*II* will be released into their inactive states. ^10^ However, an increasing body of evidence suggests that switch regions display highly flexible dynamics. ^49–52^ It is important to note that in 15% of available KRAS crystal structures S*I* exhibits a disordered conformation, in 37% S*II* is disordered, and 19% of the structures both switch regions are disordered.^51^ Structures displaying the totally open S*I* conformation were published (pdb 6M9W, 6BOF, 6QUW, and 7F0W) recently. High flexibility of S*I* and S*II* both in solution and at the membrane environments was also observed in all systems here. The chemical structures of FAR, DBD, and LIG1, including a typical snapshot of sys2 are illustrated in Fig. 2.

For the system in which KRAS4B is solvated in solution, switch regions show much higher flexibility than other systems when PM is considered, see Fig. 3 A. As predicted, switch regions of all systems exhibit a much flexible property than other moieties of CD, which is in agreement with the experimental atomic displacement parameter of switch regions, also known as B-factor that indicates the atomic fluctuations in the crystal structures,^53, 54^ indicating excellent simulation convergence. Through LIG1 binding with KRAS4B, flexibility of S*II* is largely reduced, moreover, the RMSF value of *I* of sys4 is higher than that of sys3, which agrees with published works.^11, 27, 54, 55^ Using the crystal structure as a reference, we can see that the RMSD trajectories of KRAS4B stay around 3 Å for sys1, sys2 and sys4, higher than that of sys3 (around 1.8 Å), indicating that LIG1 binding with GTP-bound KRAS4B-G12D helps it stabilize its structure and reduces its flexibility more than its GDP-bound state, see Fig. 3 C. It shows that LIG1 presents certain variant effect on KRAS4B-G12D in its GTP-bound or GDP-bound states.

**Figure 3:**
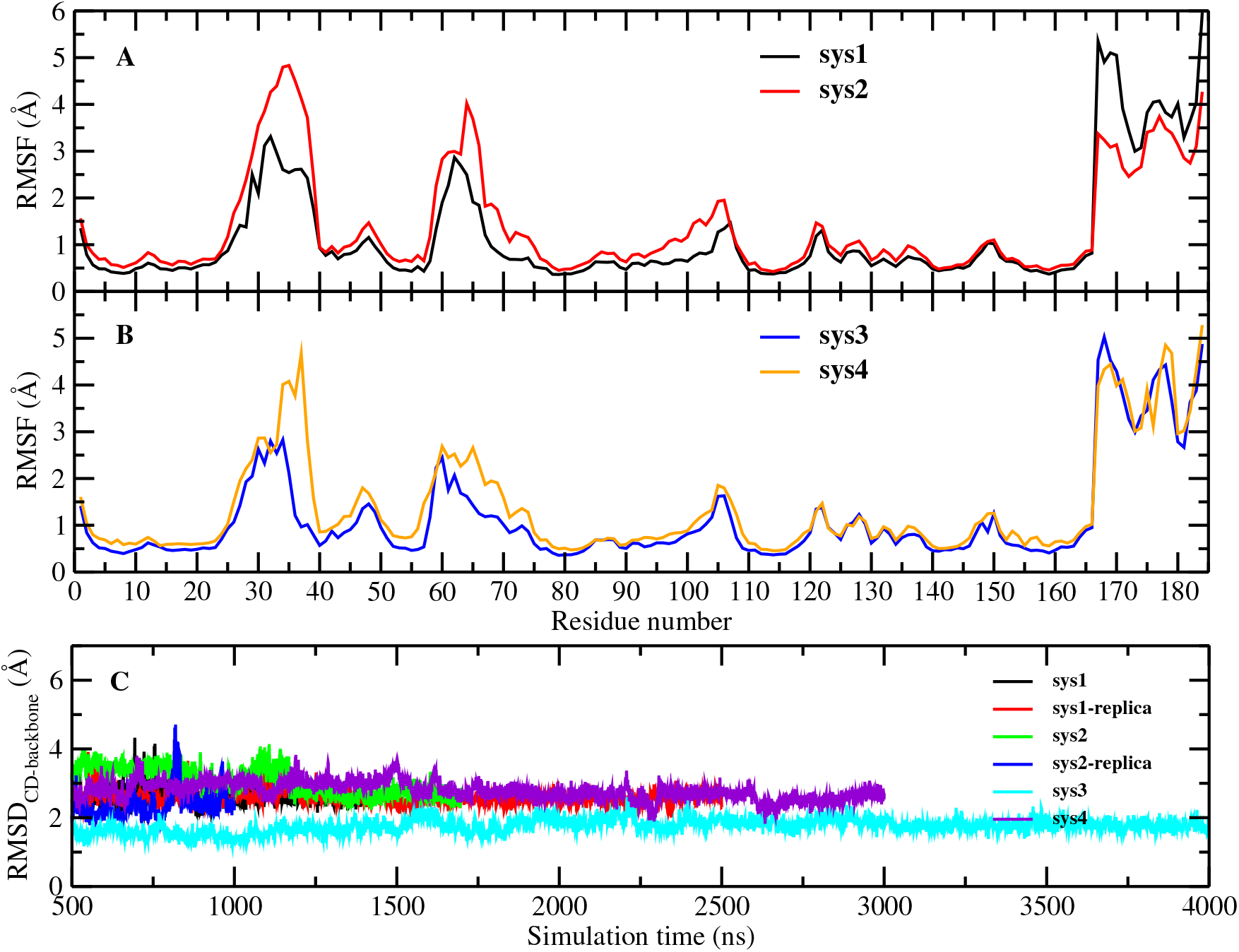
LIG1 reduces the flexibility of switch regions for both states of KRAS4B: averaged Root mean-square fluctuation (RMSF) profiles (A and B) and root mean-square difference (RMSD) of the catalytic domain of KRAS4B along the simulation time (C).

We continued to explore the corresponding atomic mechanisms. The local structure of how LIG1 establishes in the S*II* pocket in sys3 and sys4 can be analyzed by means of normalized radial distribution functions, see Fig. 4 and Fig. 5.

**Figure 4:**
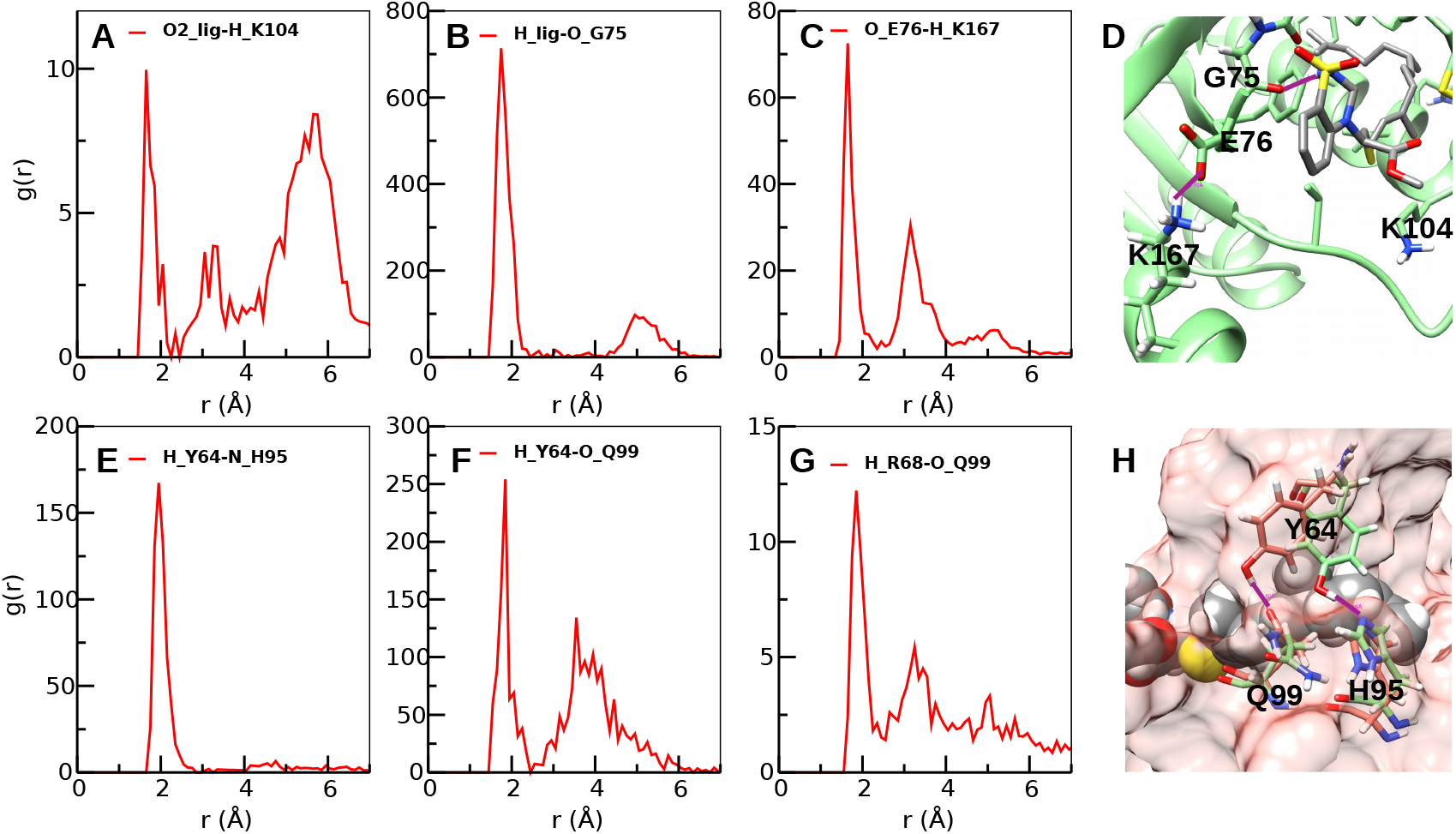
Selected averaged radial distribution functions g(r) of LIG1 with actives sites of CD of GTP-bound KRAS4B-G12D (sys3). Sites of protein residues refer to active sites of their side chains, except that “O_G75” represents its Oxygen atom of the peptide bond of the backbone. Contributions of Oxygen/Hydrogen atoms that share the negative/positive charge of side chains have been averaged. And “O1_lig”, “O2_lig”, and “H_lig” stand for active sites of LIG1 indicated in Fig. 2. D: Snapshots of typical LIG1-CD bonds. H: Snapshots of side chain of Y64 switches between residues H95 and Q99 by forming established hydrogen-bond with active nitrogen of H59 (Fig. 4E) and active Oxygen atom of Q99 (Fig. 4F) at different frames along the simulation time. Here LIG1 is shown in Van Der Waals representation for the sake of clarity. Binding sites have been highlighted in solid purple lines. D and H made with Chimera.^46^

**Figure 5:**
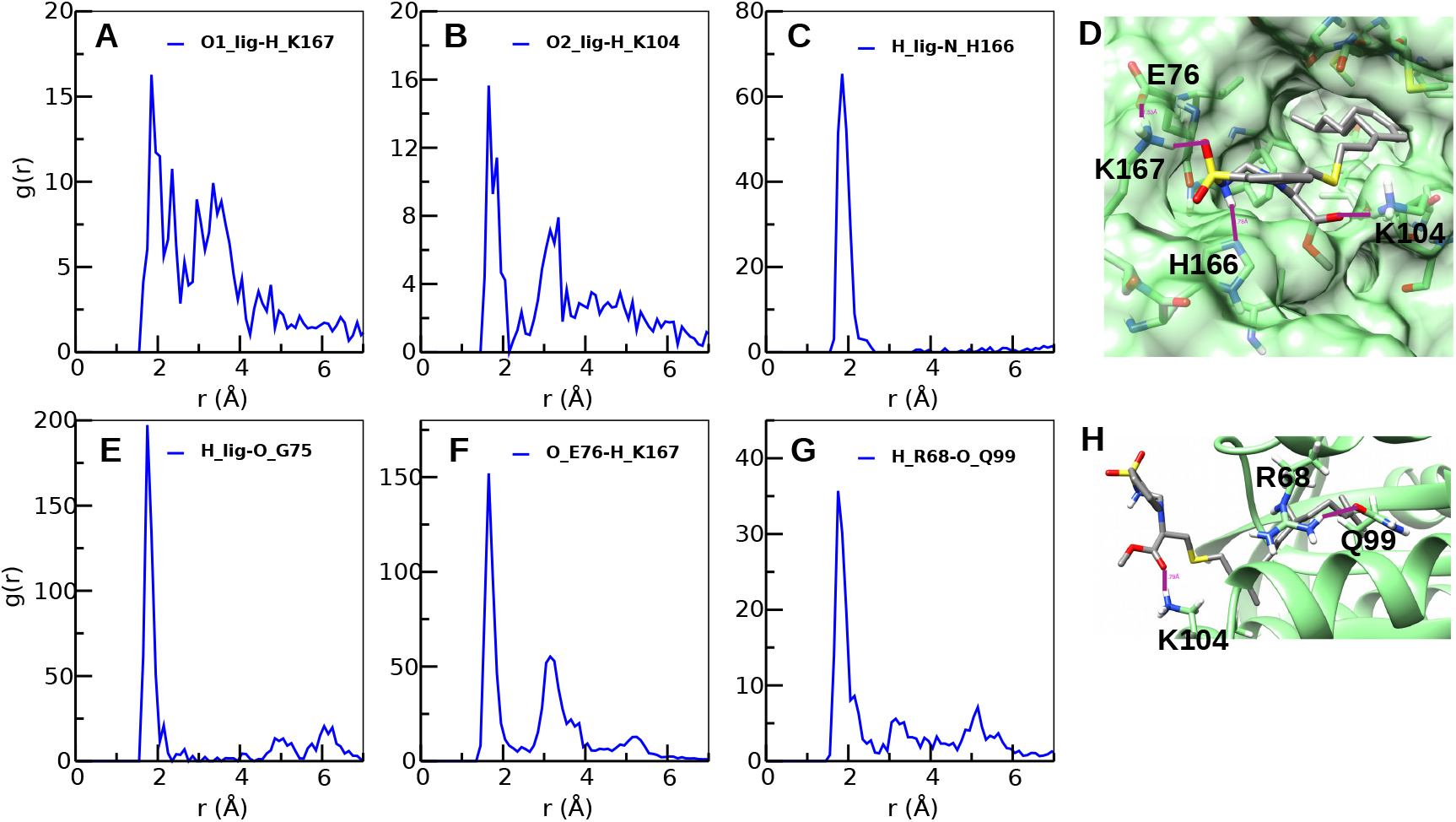
Selected averaged radial distribution functions g(r) of LIG1 with actives sites of CD of GDP-bound KRAS4B-G12D (sys4). D,H: Snapshots of typical LIG1-CD bonds. Binding sites have been highlighted in solid purple lines. D and H made with Chimera.^46^

As can be seen from Fig. 4 and Fig. 5, LIG1 is able to accommodate itself into S*II* pockets by means of hydrogen-bonds (HBs) between its DBD motif with residues G75 (Fig. 4B and Fig. 5E) and K104 (Fig. 4A and Fig. 5B) in both systems. The signature of HB in lipids and protein systems is a high peak around 1.6-2.1 Åof variable intensity.^56, 57^ Strong and long-lasting hydrogen-bond between “H_lig” and “O_G75” is observed in both cases, which helps LIG1 stably interact with S*II* region. Especially for KRAS4B-G12D in its GTP-bound state, side chain of Y64 has been found to form stable hydrogen bonding with Q99 ans H95, respectively, see Fig. 4F, G, and H. At GDP-bound state, LIG1 was found to be able to interact with residues K167 and H166 of KRAS4B through hydrogen bonds (see Fig. 5A, C, and D). Side chains of residues R68 of S*II* and Q99 of *α*-helix-3 can rotate to form a closed S*II* pocket, contributing the hydrophobic cavity shown in Fig. 5 D. These result in a much reduced flexibility in the structure of S*II* with a smaller RMSF for the GTP-bound KRAS4B than its GDP-bound state in the case of LIG1 binding into S*II* pocket, see Fig. 3 B.

### LIG1 inhibits KRAS4B-G12D in both GDP/GTP states

It has been shown that in addition to FAR insertion into the PM, the CD of KRAS4B adopts three rapidly interconverting orientation states through interactions with the anionic PM via (1) helix structures *α*-helix 3-5 on the allosteric domain while the effector-binding domain is solvent accessible (active state, OS_1_), (2) *β*-strands 1-3 on the effector-binding domain while effector-binding domain is occluded by the PM (inactive state, OS_2_), or (3) one intermediate orientation state (OS_0_).^12, 58, 59^ Proper membrane orientation state of KRAS4B directly affects its signalling activity. Here we have adopted the well-defined parameter Θ^11, 12, 59^ to describe the orientation of KRAS4B at the PM. The angle Θ is defined as the angle between the membrane normal direction and a vector running the C*_α_* atoms of the residue K5 and the residue V9 which belong to the first strand *β*1 in the structure of KRAS4B, see in Fig. 6 A.

**Figure 6:**
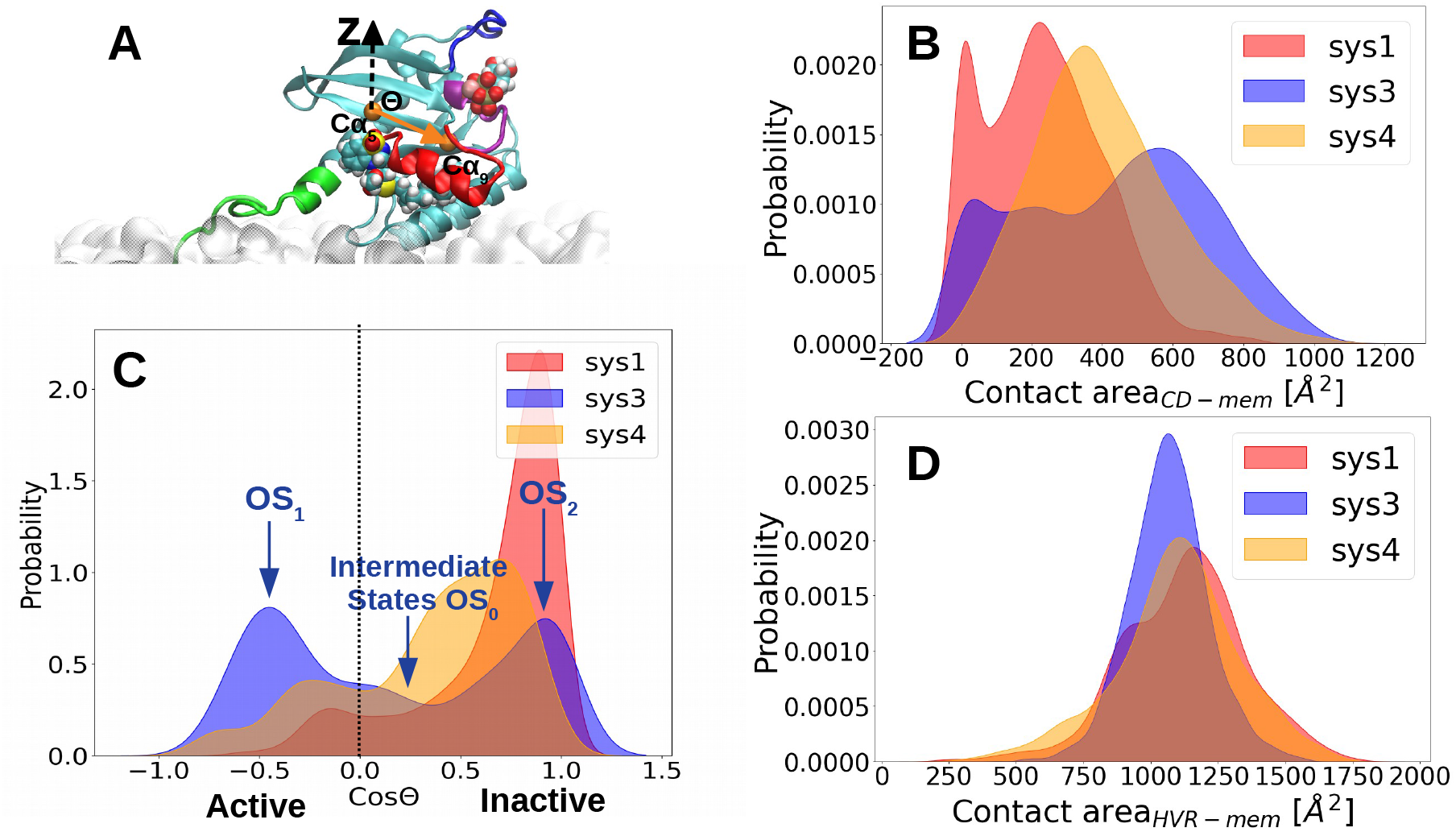
A: Definition of the orientation angle Θ, representations are the same as Fig. 2, except that the C*_α_* of residues K5 and V9 are shown in Van Der Waals in orange color. And a vertical line at CosΘ = 0 is a help for the eye, B: Distributions of contact area between CD and the PM, C: Distributions of the membrane orientations of KRAS4B in the cases of LIG1 and DBD molecules, OS_1_ and OS_2_ refer to two typical states (active and inactive, respectively) of KRAS4B observed along the trajectories of sys3, see Fig. 7 F and G, and intermediate states are indicated by OS_0_. D: Distributions of contact area between HVR and the PM.

Fig. 6 B and D shows the contact area between CD and HVR with the interface of membrane, indicating the interactions among them. The interactions between HVR and membrane show no obvious difference, showing that HVR can establish stable interactions with the anionic PM here. In general, LIG1 binding into S*II* helps KRAS4B to enhance interactions with the membrane indicated by the increased number of contact area in Fig. 6 B. The GTP-bound protein shows a larger range in the contact area of CD and membrane, indicating the widely observed exchange between different orientation states for the GTP-bound KRAS4B with LIG1 binding than the GDP-bound molecule.

Here in Fig. 6 C, we have investigated the distribution of the orientation of KRAS4B-G12D on the ionic PM under different conditions for three systems. For sys1, DBD’s presence in the system helps this mutated KRAS4B to exhibit mostly at its inactive state. However, from Figure S3 A we can see that DBD is seldom bound with CD of KRAS4B, knowing the radius of KRAS4B backbone is around 17 Å in two replica trajectories run in this work. DBD spent most of the simulation time burying itself in the interface of the PM, or being solvated in solution away from KRAS4B, distance around 75 Å in the Figure S3 A for sys1. In other words, DBD is unable to establish stable interactions with KRAS4B, which makes it hard to act as a KRAS4B-G12D inhibitor. KRAS4B of sys1 staying in its inactive state could be due to this fact, once KRAS4B tends to stay in its current metastate (inactive state in this case) in the time scale of MD simulations considered here. It is harder to transit between orientation states (OS_2_-OS_0_-OS_1_) while anchoring to the PM compared to KRAS4B with LIG1 bound to its S*II* pocket of sys3. It reveals here that with LIG1 binding into KRAS4B, the G12D mutant was populated averagely in its three states (active, intermediate, and inactive states), showing a much less cancerous property. Hence, it shows that LIG1 helps the GTP-bound KRAS4B to exchange more easily between its orientation states on PM which makes it less “cancerous”, knowing that KRAS4B-G12D without LIG1 tends to stay mostly in its active orientation state. In 2016, wild-type GTP-bound KRAS4B has been reported to exist mostly in its active state.^60^ LIG1 binding into S*II* pocket of GTP-bound mutated KRAS4B may trigger a population shift toward the inactive state hindering the effector-binding site and switching the catalytic domain orientation of KRAS4B. Meanwhile, LIG1 reveals to be able to lock GDP-bound KRAS4B-G12D to exhibit the inactive orientation at the PM while effectively hiding the effector-binding interfaces, inhibiting the guanine nucleotide exchange, resulting in a mostly GDP-bound state such that the following signaling pathways are inhibited, under circumstance of sys3 and sys4 sharing the same initial orientation and conformation in the structure.

### KRAS4B anchoring to PM is crucial to fulfill its physiological function

To observe the effects of DBD and LIG1 on protein dynamics, we calculated the cross-correlation of the atomic fluctuations obtained from their MD trajectories. The transmission of the allosteric signal within the protein shall couple motion between active site residues.^61^

From Fig. 7, a similar pattern of positive and negative correlations in motion were observed for systems of KRAS4B anchoring into PM. DBD is considered in sys1, positive correlated motion is still observed in its effector-binding domain, which is marked in blue box in Fig. 7 A, even the mutated KRAS4B presents in the accumulated inactive orientation states, see Fig. 6 C. We have observed that LIG1 helps GTP-bound KRAS4B-G12D to present a balanced distribution in the active (OS_1_) and inactive (OS_2_) states, see Fig. 6 C. In Fig. 7 C and D, we can observe that LIG1 directly binding with KRAS4B effectively diminished correlations in the effector-binding motif, whereas the increased negative correlated motion between the helix3 (res 87-104) and the effector-binding domain is observed for GDP-bound protein. However, when solvated in the aqueous environment, it is obvious to notice that both negative and positive correlations in motion have been lost, which agrees with Jang’s work,^60^ indicating a decreased cross-talk in the structure of KRAS4B. This may indicate that PM is crucial for KRAS4B’s dynamics in order to transmit signal across its structure. We have also taken a close investigation on the cross-correlation of switch regions, see Figure S4. We can observe that strong correlations in motion in the interswitch regions are conserved in all systmes in PM environment. Effector-binding regions which are important for effector recruitment have lost correlations in motion for KRAS4B in solution environment. Only for KRAS4B at the bilayer with proper orientation positive or negative correlated motions can be observed. Our results are in good agreement with the published work.^60^ Together, KRAS4B anchoring to PM is crucial to fulfill its physiological function. Therefore LIG1 binding into S*II* pocket of mutated KRAS4B inhibit its proper biological function by interfering the corresponding cross-talk throughout its structure.

**Figure 7:**
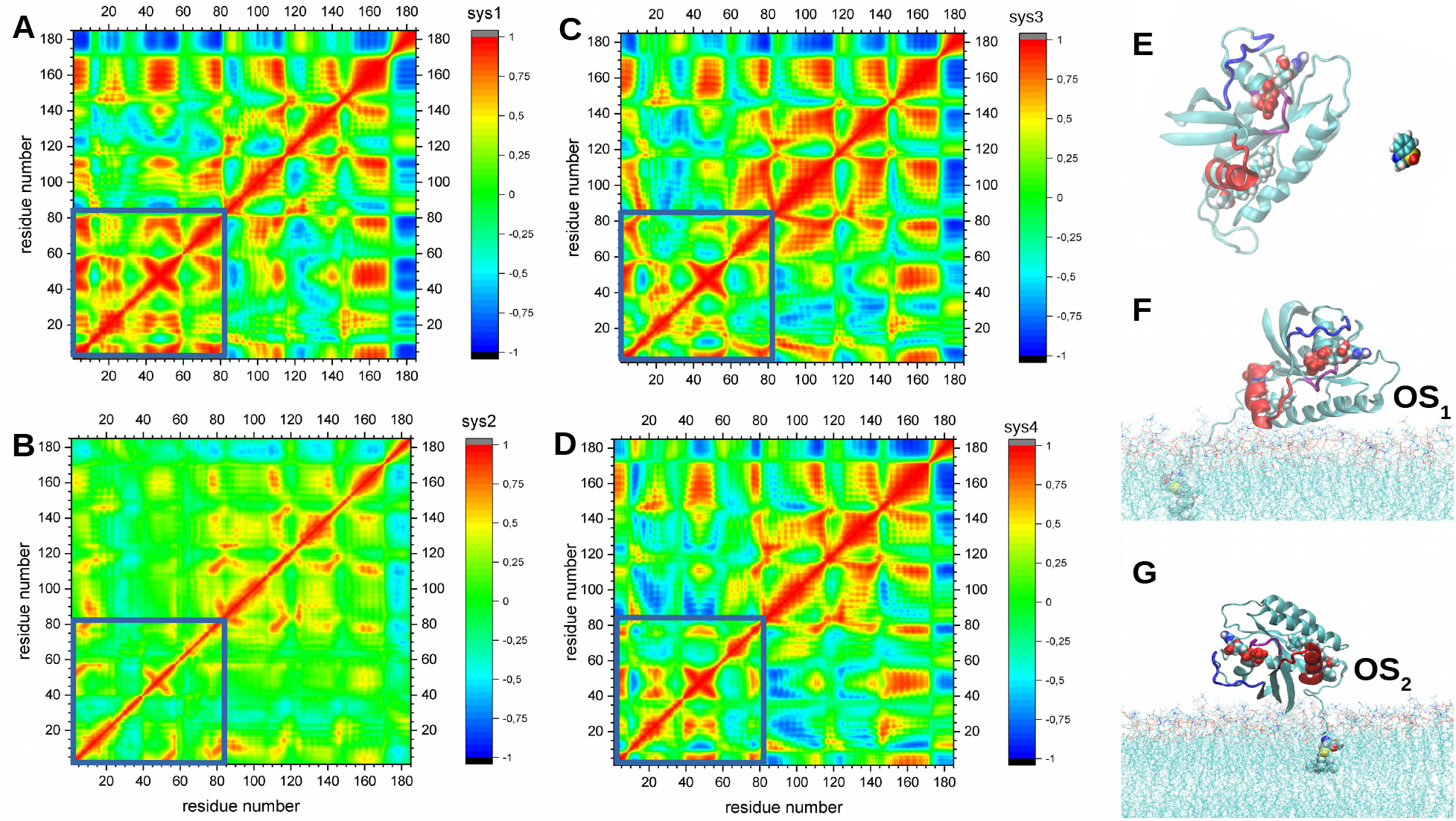
A, B, C, and D: Comparison of correlation analysis of the motion during the final 1000 ns MD simulation of different systems studied. Highly (anti)correlated motion are orange or red (blue). Correlations within effector-binding domain are marked in blue boxes. E: typical configuration of KRAS4B in the solution environment of sys2, DBD is observed to be away from KRAS4B for the most of trajectories. F and G refer to two typical orientations of KRAS4B at the PM, snapshots are taken from different frames of sys3 and made with VMD.^62^

### LIG1 acts as the KRAS4B competitor to bind with PDE***δ***

Signal transduction of KRAS4B is strongly linked to its enrichment at the PM.^63^ The solubilization factor PDE*δ* has been reported to play the role in sequestering KRAS4B from the cytosol by binding the FAR, thereby, enhancing its diffusion throughout the cell, then releasing KRAS4B to bind with the PM.^64, 65^ Inhibition or knock down of PDE*δ* has been proved itself to be essential to control KRAS4B’s enrichment at the membrane which is crucial for growth factor signaling output. LIG1 shares the same FAR motif as full-length KRAS4B, so we have investigated whether LIG1 can act as the PDE*δ* inhibitor to impair KRAS4B enrichment at PM. The crystal structure (PDB 5TAR) of PDE*δ* was taken as the initial structure of the complex of PDE*δ*-LIG1, and the interactions of FAR moiety and PDE*δ* of PDB 5TAR were preserved for LIG1 and PDE*δ*.

In Fig. 8, the averaged value of RMSD profile is of around 1.9 Å, a low value together with the corresponding RMSF profile, showing a steady structure of PDE*δ* due to the antiparallel beta strands in its structure. Fig. 8 D shows clearly that LIG1 accommodates itself steadily in the cavity formed by the hydrophobic side chains of related residues. Meanwhile, LIG1 can also establish hydrogen bonds with side chains T149 and A112 of PDE*δ* through its active sites of the inimo and sulfonyl groups of the DBD moiety, indicating high binding affinity between LIG1 and PDE*δ* protein. Taken together, LIG1 might inhibit the cancerous pathways that KRAS4B gets involved in both ways.

**Figure 8:**
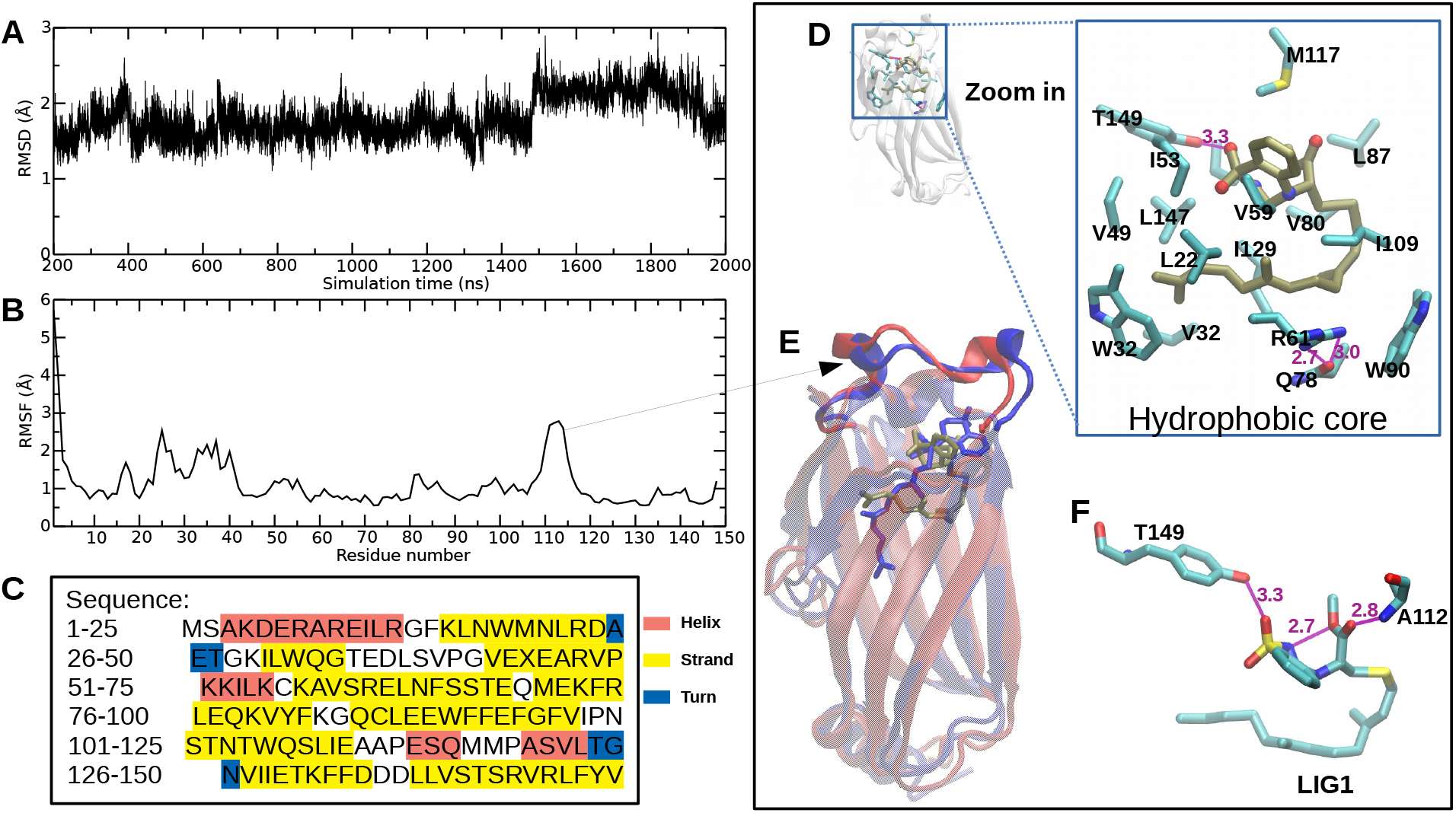
A, B and C: RMSD trajectories, RMSF plot and sequence of PDE*δ*, D: detailed hydrophobic interactions of LIG1 (tan sticks) and PDE*δ*, E: superimposed structure of PDE*δ* of the crystal structure (blue) and last (red) frame of production runs for 2000 ns simulation time showing no dramatic changes in the structure of PDE*δ*. F: hydrogen bonding between LIG1 and T149 and A112 of PDE*δ*, H atoms of T149, A112, and LIG1 not shown for the sake of clarity.

## Conclusion

Molecular dynamics simulations have become a very valuable computational tool for drug discovery. Once a reliable force field is assumed, MD have the advantage over wet-lab experiments that we can monitor dynamics on real time and explore the stability and suitability of potential drugs. Inhibitors with high potency of intracellular biologics to reduce the required level of intracellular delivery, and consequently to facilitate the development of “cell-penetrating” biologics targeting intracellular proteins are in urgent need. Drugs using lipid-based formulation usually conserve good potential in cell permeability properties. In this work we describe a designed drug LIG1 using lipid-based formulation that directly interacts with KRAS4B-G12D in both nucleotide states and its regulator PDE*δ* through hydrophobic force by its FAR tail and hydrogen bonds by the DBD motif to inhibit KRAS4B’s abnormal signaling activities. Our findings also suggest that KRAS4B anchoring to PM is crucial to conserve its intramolecular communication between different residues/domains to fulfill its physiological function, hence, considering PM during its corresponding drug discovery for KRAS4B and other membrane oncogene proteins is essential and that has been neglected for a long time.

LIG1 reveals to be able to lock GDP-bound KRAS4B-G12D to exhibit the inactive orientations at the PM while effectively hiding the effector-binding interfaces, which leads to inhibit the guanine nucleotide exchange. Meanwhile, LIG1 reveals to trigger a population shift to exist in its inactive states for GTP-bound congogenic KRAS4B. Therefore, the following signaling pathways in which GTP-bound KRAS4B involves are inhibited. Our study shows that LIG1 directly targets PDE*δ* which inhibits the KRAS4B-PDE*δ* interaction and impairs KRAS4B enrichment at the membrane, thereby suppresses the *in vitro* and *in vivo* proliferation of KRAS-dependent cancer cells, further deepening our understanding of drug-KRAS4B recognition. It would be interesting to examine the molecular mechanisms/interactions between LIG1 and its derivatives to target different oncogenic KRAS4B mutants, or other RAS species, since up to date lipid-based drugs have not been considered to target RAS mutants. Beyond the main-stream strategy of designing anti-cancer drugs, our work could be exploited by clinical strategies for anti-cancer drug design and provide a novel opportunity to suppress KRAS4B-related tumour growth.

## Methods

### MD simulation details

Five model systems were constructed to systematically examine that how the drug (LIG1) designed here can affect KRAS4B-G12D’s orientation and corresponding signalling pathways. Along with this work, we have also tested the possibility of a small drug DBD works as the warhead to target the KRAS4B-G12D, with the setup characteristics for all simulations listed in Table 1.

Anionic cell membranes constituted by DOPC (56%), DOPS (14%) and cholesterol (30%) were considered for all systems. These membrane systems contains a total of 308 lipid molecules fully solvated by TIP3P water molecules in potassium chloride solution at the human body concentration (0.15 M). DBD molecule was localized near KRAS4B at the same side of PM while solvated by water molecules and KRAS4B was initially set anchoring to membrane at the beginning of simulations in membrane systems, see Figure S5. For water system, DBD was placed around 30 Å away from KRAS4B along the Y axis. All MD inputs were generated using CHARMM-GUI membrane builder^66, 67^ and the CHARMM36m force field (FF)^68^ was adopted for lipid-lipid and lipid-protein interactions. Crystal structure of KRAS4B with partially disordered hypervariable region (pdb 5TB5), GTP (pdb 5VQ2) and GDP (pdb 5xco) were used to generate full length GTP-bound/GDP-bound KRAS4B proteins, and parameters of DBD were directly adopted from the charmm36m FF topology files. The crystal structure (pdb 5TAR) was used to generate the full-lenth PDE*δ*, interactions between FAR tail and PDE*δ* were conserved in the crystal structure, see in Fig. 8 E. All pdb files were downloaded from RCSB PDB Protein Data Bank.^69^ The N-termini and C-termini of KRAS4B were set as NH^+^ and COCH_3_ groups, respectively. All-atom MD simulations were conducted using the AMBER20.^70^ All systems were energy minimized for 20000 steps followed by five 250 ps equilibrium runs while gradually reducing the harmonic constraints on the systems. Minimization steps for replicated systems of sys1 and sys2 were set to be 5000 and 10000 steps, rather than 2000 steps, to generate independent production trajectories. Production runs were performed with NPT ensemble, Langevin dynamics with the collision frequency 1.0 ps*^−^*^1^ was used for temperature regulation (300 K) and Monte Carlo barostat was used for the pressure regulation (1 bar), respectively. Semiisotropic pressure scaling with constant surface tension for xy plane is used in statistical ensembles for simulating liquid interfaces. The time step was set to be 4 fs and the frames were saved by every 25000 steps for analysis. The particle mesh Ewald method was used to calculate the electrostatic interaction, and the van der Waals interactions were calculated using a cutoff of 12 Å. Periodic boundary conditions in three directions of space have been taken.

### MD simulation analysis

The root mean squared deviation (RMSD), root mean square fluctuation (RMSF), and contact area through monitoring the solvent-accessible surface area (SASA) along the trajectories for different structures are calculated using Tcl scripts by VMD.^62^ Contact map analysis was constructed by software ConAn^71^ to analysis the residue-residue interaction. We define a contact as the distance between two heavy atoms being with a distance cutoff. The contact cutoff for two carbon atoms is set as 6 Å, the other heavy atoms for 4.5 Å. The normalized covariance (correlation) of the MD simulations was performed by using CARMA,^72^ which gave rise to a residue-residue correlation map. A set of nodes in the correlation map with connecting edges, which in this case amino acid nodes were centered C*_α_* atoms. Here we define that the edges between nodes whose residues are within 4.5 Å for at least 75% of the frames analyzed. Intra-molecular correlations between residues were measured by normalizing the cross correlation matrix of atomic fluctuations over the length of the simulation.^73^ If two residues move in the same (opposite) direction in most the frames, the motion is considered as (anti-)correlated, and the correlation value is close to −1 or 1. If the correlation value is close to zero, these two residues are considered uncorrelated.

## Data Availability

Input files for the production runs of MD simulations, initial coordinates of five systems studied in this work can be found in our GitHub repository at https://github.com/soft-matter-theory-at-icmab-csic/anticancer-drug-KRAS4B-G12D. We also provide some scripts of the MD analysis.

## Acknowledgement

We thank financial support by the Margarita Salas grant which is funded by the European Union - NextGenerationEU awarded to the Universitat Politecnica de Catalunya (Huixia Lu). We also thank financial support by Grant PID2021-124297NB-C32 and PID2021-124297NB-C33 funded by MCIN/AEI/10.13039/501100011033, “ERDF A way of making Europe” and given by the “European Union NextGenerationEU/PRTR”. We thank the CESGA supercomputing center for computer time and technical support at the Finisterrae III supercomputer. Molecular graphics made with UCSF Chimera, developed by the Resource for Biocomputing, Visualization, and Informatics at the University of California, San Francisco, with support from NIH P41-GM103311. Zheyao Hu is a Ph.D. fellow from the China Scholarship Council (grant 202006230070).

## Author Contributions

H.L. and J.M. designed the study. H.L. performed the simulations, analyzed the data, and generated the figures. All authors took part in writing, discussing and revising the manuscript.

## Supplementary information for: Discovery of a novel drug using lipid-based formulation targeting G12D-mutated KRAS4B through non-covalent bonds

Recently reported warheads include *α*,*β*-unsaturated carbonyl moieties such as AMG510^1^ and MRTX849,^2^ electrophilic warheads such as heteroaromatic motifs, sulfur(*V I*) motifs, terminal alkynes motifs, etc. see Figure S1. Examples of chemical structures of FDA approved drugs using lipid-based formulations are listed in Figure S2. Figure S3 records the corresponding distance profiles of mass of center of interest. In Figure S3 B, we can observe that FAR binds into S*II* of KRAS4B in two independent trajectories, showing a great tendency of FAR serving as the binding hook of the newly designed drug reported here. And Figure S3 D shows that LIG1 is able to bind with KRAS4B-G12D in both nucleotide states. Figure S4 exhibits the two-dimensional dynamics cross-correlation map of the residues motions of switch regions. The initial configurations of KRAS4B-related systems are illustrated in Figure S5.

**Figure S1:**
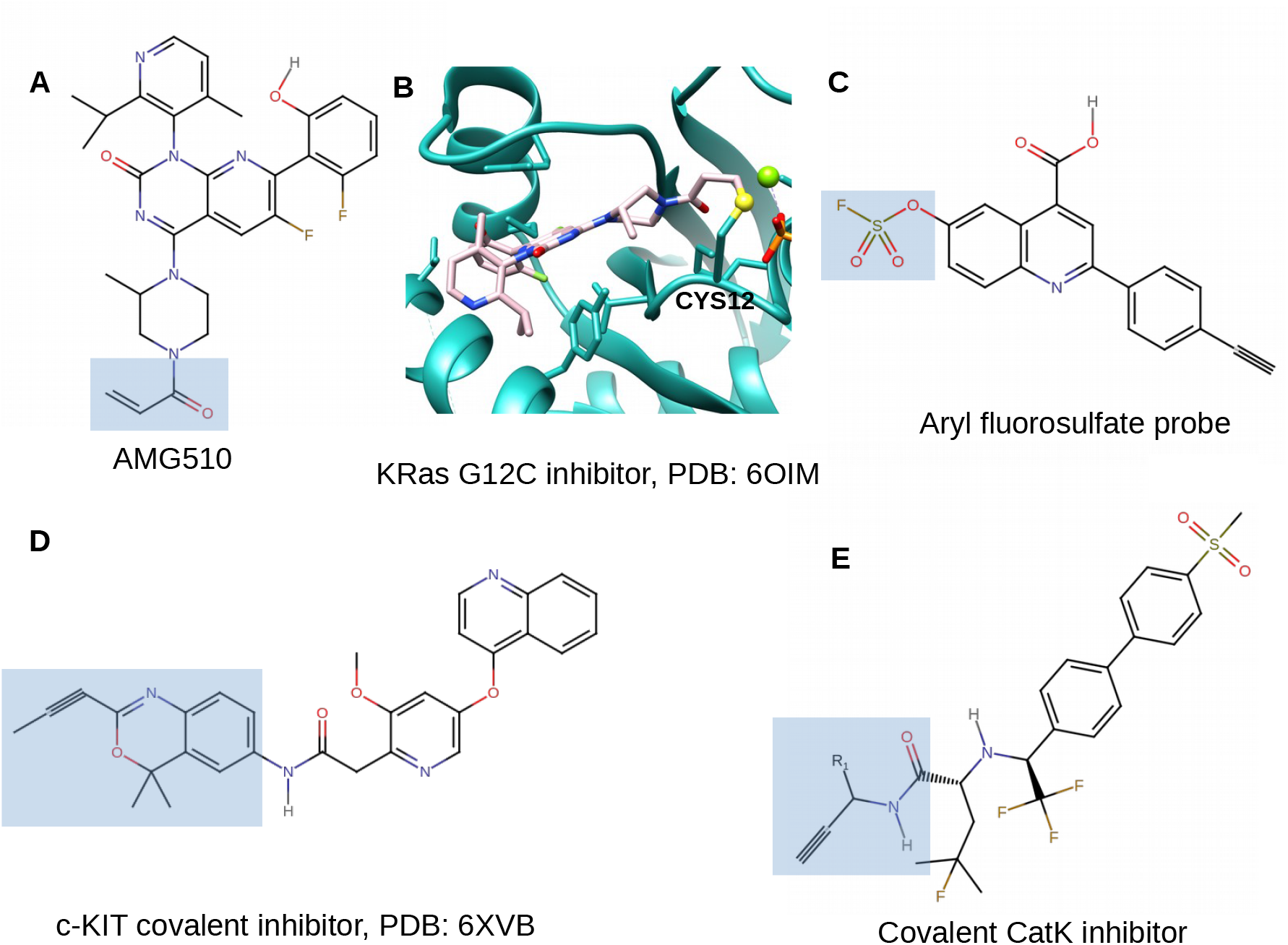
Examples of different warheads marked inside blue boxes: A. *α*,*β*-unsaturated carbonyl warhead: Chemical structure of AMG510, B. X-ray crystal structure of AMG510 bound to CYS12 of KRAS-G12C, AMG510 shown in pink color, C. Chemical structure of one sulfur(*V I*) warhead,^3^ D. Chemical structure of one heteroaromatic warhead,^4^ and E. Chemical structure of the terminal alkyne moiety-based CatK covalent inhibitor.^5^

**Figure S2:**
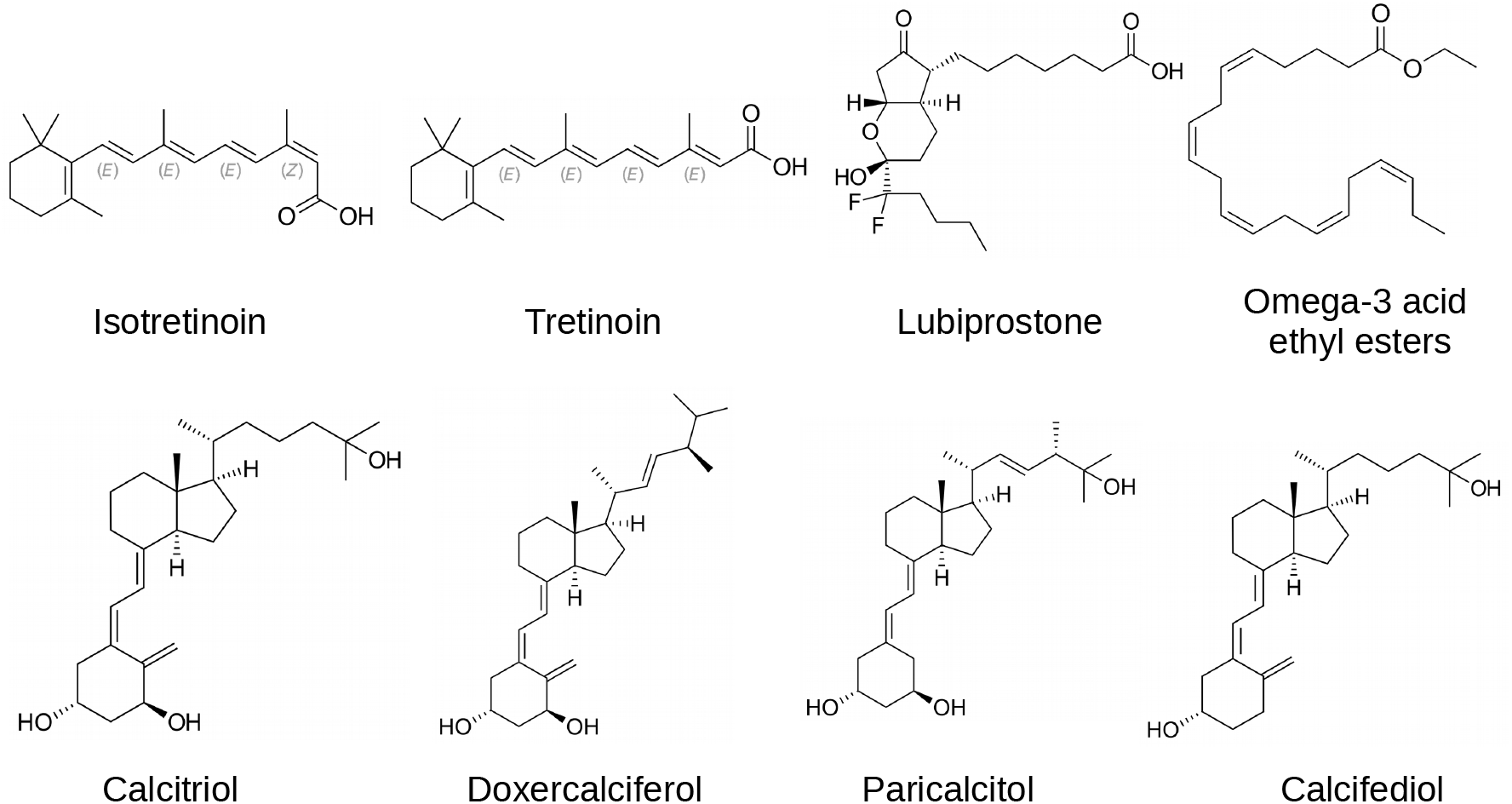
Chemical structures of some FDA approved drugs using lipid-based formulations. ^6^

**Figure S3:**
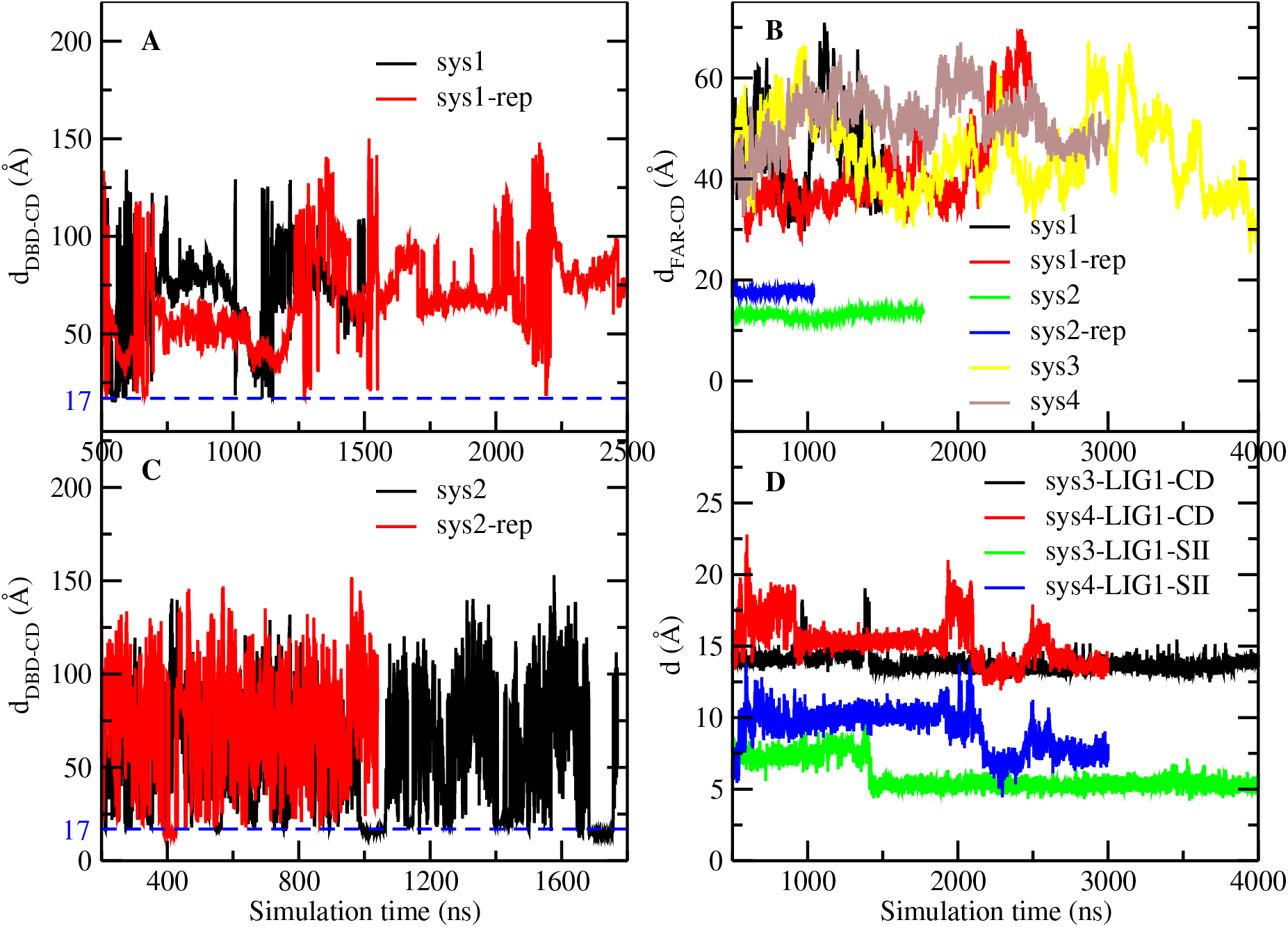
Selected distance profiles of mass of center along with the simulation time.

**Figure S4:**
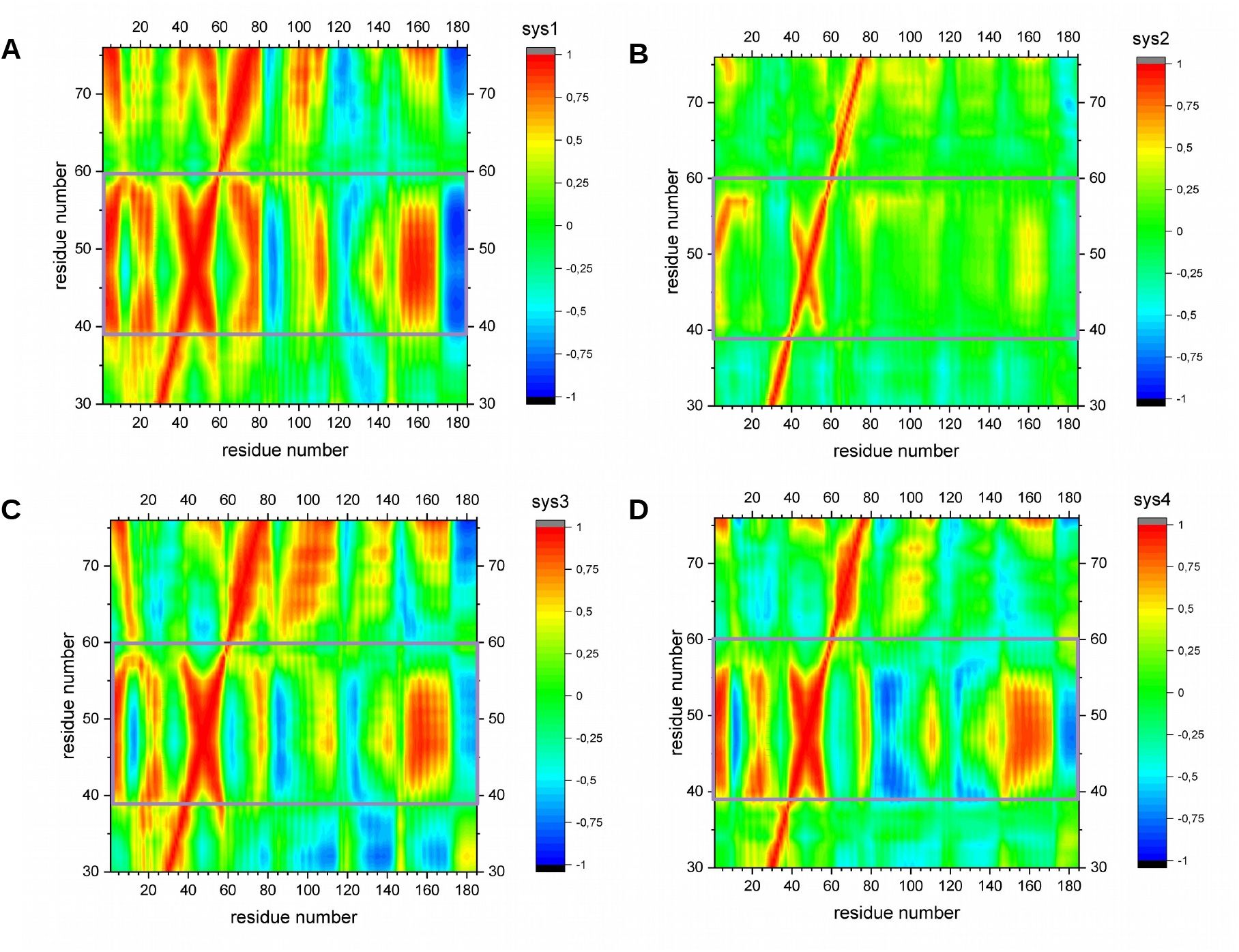
Two-dimensional dynamics cross-correlation map of the residues motions of the switch regions for KRAS4B systems studied here.

**Figure S5:**
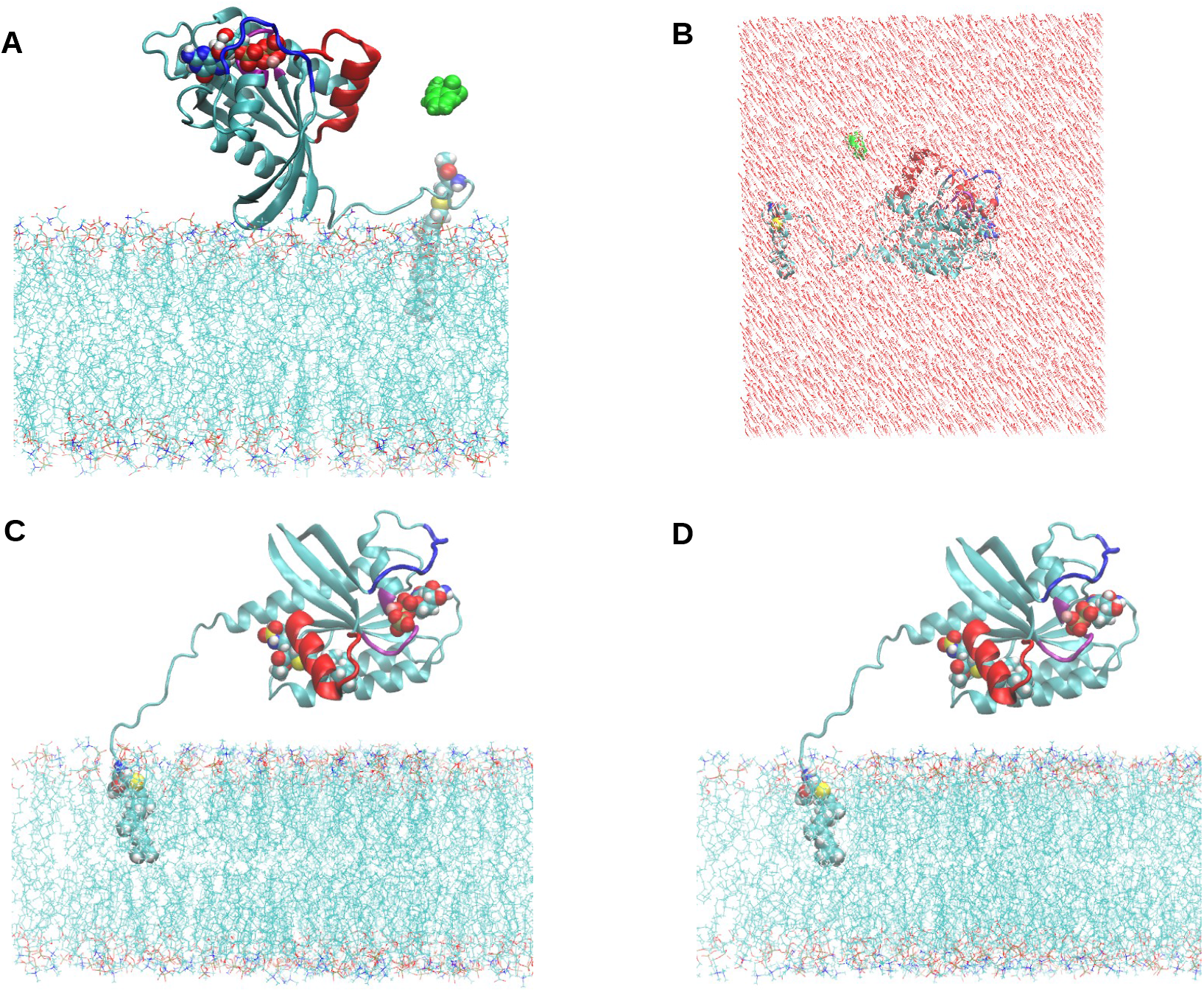
Initial Configurations of KRAS4B systems studied in this work: A(sys1), B(sys2), C(sys3), and D(sys4). Water molecules and ions are not shown for systems in which membrane is included for the sake of clarity. Herein, PM is shown in line, KRAS4B shown in NewCartoon, and DBD molecule (green), Mg ion (pink), GTP/GDP, and LIG1 are in depicted using VDW style. S*I*, S*II*, and P loop are shown in NewCartoon representation in color blue, red, and purple, respectively.

